# Concurrent processes set *E. coli* cell division

**DOI:** 10.1101/301671

**Authors:** Gabriele Micali, Jacopo Grilli, Matteo Osella, Marco Cosentino Lagomarsino

**Author notes:** Equal Contribution (Gabriele Micali, Jacopo Grilli).

## Abstract

A cell can divide only upon completion of chromosome segregation, or its daughters would lose genetic material [1, 2]. In *E*. *coli* bacteria, the prevalent view is that cells divide a fixed amount of time after they start to copy the chromosomes [3, 4], and a known pathway prevents cells from dividing if the chromosomes interfere with the cytokinesis machinery [5]. However, whether completion of segregation is typically the bottleneck process for the decision to divide has never been stringently tested on single cells. We show how key trends in single-cell data lead to challenge the classic idea of replication-segregation limiting cell division. Instead, the data agree with a model where *two* concurrent processes (setting replication initiation and inter-division time) set cell division on competing time scales. During each cell cycle, division is set by the slowest process (an “AND” gate). The concept of transitions between cell-cycle stages as decisional processes integrating multiple inputs instead of cascading from orchestrated steps can affect the way we think of the cell cycle in general.

Dynamic single-cell data revived the classic debate on the determinants of cell division, but the recent literature is fragmented into different and contrasting models [3]. Most studies take the classic view that a fixed “*C*+*D* period”, comprising a “C period” to copy the genome and a “D period”, needed to complete segregation and running from termination to cell division (Fig. 1a) is rate-limiting for cell division (Fig. 1b), but these studies do not agree on the underlying mechanisms [6, 7]. A recent study by Harris and Theriot formulates the opposite hypothesis [8]. The main assumption of this alternative view is that the rate-limiting process for cell division is instead the completion of the septum (Fig. 1b), and consequently chromosome segregation is *never* rate-limiting for cell division.

**FIG. 1.**
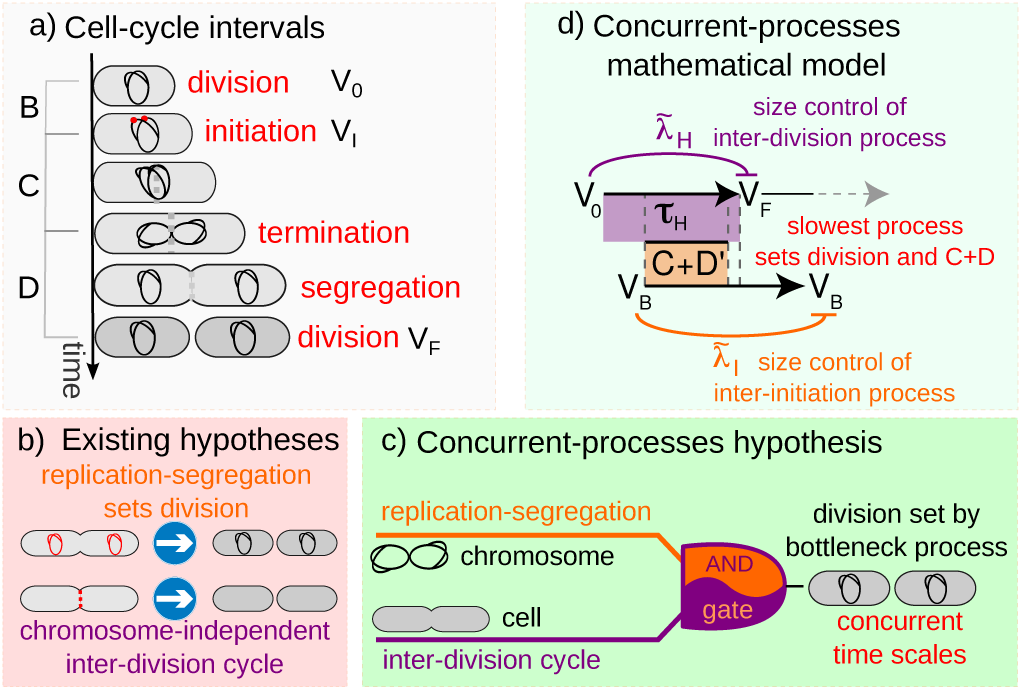
The concurrent-processes hypothesis. a) Explanation of the replication-related cell-cycle intervals in *E*. *coli*. Replication initiation occurs after a “*B* period” followed by the *C* (replication) and *D* (termination to division) periods, the *B* and *C* + *D* period can be measured in single cells by proxies of replication initiation [6, 11]. b) Classically, replication-segregation is believed to be rate-limiting for cell division; a recent hypothesis by Harris and Theriot [8] states that the rate-limiting process might be instead the formation of the septum. c) Our concurrent-processes hypothesis states that cell division is the result of the slowest between an interdivision cycle (setting division when, e.g., the septum machinery is ready) and a replication-related processes (setting division when replication-segregation is complete). Hence, the circuit is analogous to an AND gate. d) Scheme of the mathematical model.

More specifically, the authors provide evidence that surface synthesis rate is proportional to volume, and they propose a model where division is set by a threshold surface to synthesize the septum [8]. This model recapitulates the empirical size-control strategy followed by these cells, whereby the added size is nearly constant, regardless of initial size [8–10] (the so-called “adder” behavior). Clearly, with such contrasting results the question of which process drives cell division becomes pressing (Fig 1b). Our central finding is that *both processes* concur to set cell division (Fig 1cd).

To support this point we started our analysis from the available experimental data, and we concentrated on two underrated correlation patterns measured recently for the *C*+*D* period. These patterns are shown in Fig 2ab. First, we find that the growth of the cell during the *C* + *D* period, quantified by the log ratio of the division volume and the initiation volume, is anticorrelated with cell size at initiation (Fig 2a). This pattern is very consistent across strains, measurement methods, and conditions, but overlooked by current models assuming that replication is the bottleneck process [6, 7]. Second, we find that the duration of the *C* + *D* period is clearly anticorrelated with the growth rate of individual cells, with a near-inverse relationship (Fig 2b), as reported by Wallden and coworkers [6].

**FIG. 2.**
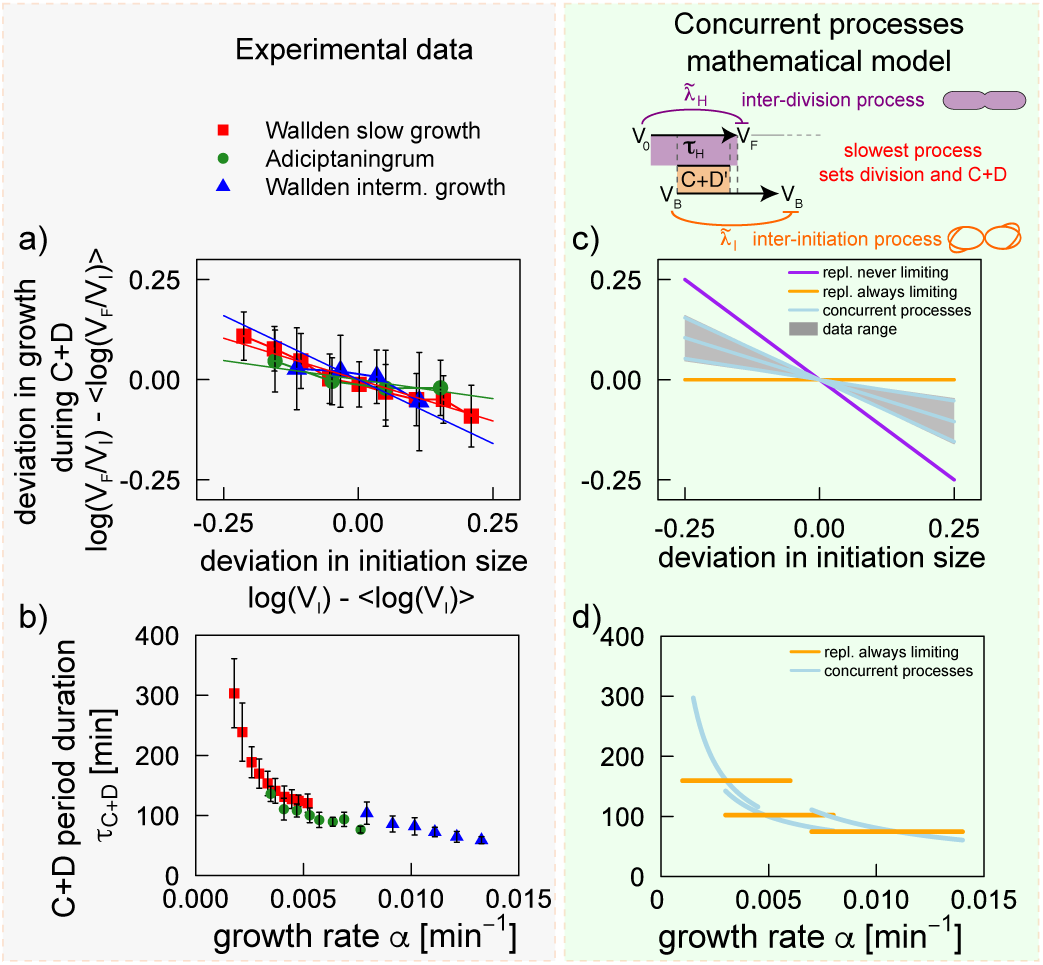
A concurrent-processes model explains the correlation patterns for the *C* + *D* period. Panels a and b plot the unexplained patterns for the *C* + *D* period. a) Cell growth during the *C* + *D* period, quantified by the logarithmic ratio between final and initial volume (*y* axis, binned averages), anti-correlates with cell size at initiation (*x* axis). The plot is centered by the mean values of the *x*– and *y*-axis variables to compare different data sets. Solid lines are linear fits. b) The duration of the *C* + *D* period (*y* axis, binned averages) anticorrelates with the growth rate of individual cells (*x* axis), with a near-inverse pattern. c) The correlation between size at initiation and growth during the *C* + *D* period (panel a) attains the observed intermediate slopes (gray shaded area) in the concurrent-processes hypothesis (cyan lines, see methods for parameters), but not if one assumes that replication is always (orange line) or never limiting (purple line) cell division. Solid lines correspond to theoretical predictions and the shaded area to the range of slopes allowed by the plots in panel a. d) The anticorrelation between *C* + *D* period duration and growth rate of individual cells (panel b) is absent if replication is always bottleneck (orange line). Instead, the concurrent-processes hypothesis captures experimental trends quantitatively (cyan lines have the same parameters as in panel b, see methods). Solid lines correspond to theoretical predictions. Data from refs [6, 11].

These patterns are not compatible with the classic assumption of replication driving division: there are no available justifications for such correlations between timing and either growth rate or cell size. If replication is always the bottleneck process, division happens on average at a fixed time after initiation of replication. Assuming that this fixed time is not coupled to cell size or growth rate, both plots in Fig. 2ab would show no correlation (Fig. 2cd, orange lines).

In contrast, the *C* + *D* patterns emerge naturally from a model in which the division event is set by two concurrent processes. In a concurrent processes framework, the *C* + *D* period is the juxtaposition of two processes: the time by which the replication-related process is ready to divide, defined as *C* + *D*′, and the time by which the inter-division process (e.g., the septum) is completed. The measurable *C* + *D* is the time for the cell to really divide and is set at the single-cell level by the slowest between these two processes. Whenever division is set by the inter-division process, the cells that initiate replication at a larger size will grow less during the *C* + *D* period (because in this case the final size ignores the initiation size), which gives the pattern of Fig. 2a. Equally, each time the inter-division process is slowest, the *D* period will tend to be shorter for cells growing faster than average due to the size-based inter-division control. For example if the inter-division process adds an average constant size [8, 9], this target size will be reached faster by single cells growing faster than average, which will decrease the duration of their *C* + *D* period. This yields the pattern shown in Fig 2b.

To go beyond these qualitative considerations, and to produce testable quantitative predictions on the assumption of concurrent processes setting cell division, we formulated and solved a mathematical model based on this principle (see methods). In this model, two size-dependent processes setting the inter-initiation time and the inter-division time run in parallel, and division follows the slowest process (as in an AND gate) between completion of the inter-division process and completion of a *C* + *D*′ period after initiation (Fig. 1d). We solve this model both analytically and by numerical simulation. Fig. 2cd shows the predicted patterns of the *C* + *D* period (light blue lines). The model reproduces both correlation patterns shown in Fig. 2ab. Fig 2c also shows that when replication is limiting, both slopes are flat (orange lines in Fig. 2cd), clearly indicating that the classic framework is too restrictive. Instead, models assuming that replication is never limiting [8] may reproduce the trends in Fig. 2ac, but the slope is quantitatively too strong for the data in Fig 2a. The data reside in the intermediate regime, where competition between the two processes setting cell division is relevant (see also ref. [12]). Overall, the concurrent-processes assumption explains the patterns for the *C*+*D* period naturally from the conceptually simple assumption of competing time scales between the different processes that set cell division.

Importantly, the fact that the model captures the data does not depend on a specific parameter set, but emerges from the hypothesis of concurrent processes. This point can be shown by direct analytical estimates of the patterns in Fig. 2 (see methods for details). Our model (see Fig. 1d and methods) is specified, together by the two size control parameters λ̃_*I*_ and λ̃_H_, (ranging from 0, strong control, to 1, no control), by the characteristic initiation and division sizes encoded by the two concurrent processes, and the intrinsic cell-to-cell stochasticity of their duration (see methods). The values of these parameters affect the probability *p_H_* that the inter-division process is rate-limiting, which is the only relevant emergent parameter of the model. For any parameter set, the plots in Fig. 2 depend on *p_H_* only: they deviate from constancy if replication is not always limiting, i.e. when *p_H_* deviates from 0, and the case of replication never limiting (*p_H_* = 1) appears quantitatively too extreme to fit the data. In the data, we estimate from the slopes that *p_H_* is between 0.2 and 0.7 (see methods), well within the parameter region in which there is actual competition between the two processes. Thus, regardless of parameter values and details, only competing time scales reproduce efficiently the data.

We found that similar conclusions apply to other testable predictions of the model, beyond the *C* + *D* period. First, the model predicts that competition between concurrent processes should affect the relation between the growth in the *B* and *C* + *D* periods. To test this, we employ the method introduced by Chandler-Brown and coworkers in yeast [13], comparing growth (measured here by log ratio of final to initial volume) in the two periods. If replication is never limiting, fluctuations in the initiation size make the *B* period longer and the *C* + *D* period shorter, affecting in opposite directions growth in the *B* and *C* + *D* periods, which become anti-correlated. If replication is always limiting, growth in the *C* + *D* period should instead be independent from growth in the *B* period. Our results show that the concurrent-processes model predicts an intermediate situation between these two, which is indeed where all the data are found (Fig. 3ab). Second, competition between concurrent processes should also affect measurable patterns for the inter-division cycle, where abundant data are available. Motivated from the trend in Fig 2b, we focused on the anti-correlation reported between inter-division time and individual cell growth rates [14]. Our results (Fig. 3cd) show that the model captures the decreasing trend of this correlation with average growth rate. Once again, rather than the specific numerical values of the parameters, we find that the data take intermediate values between the two extreme cases of replication always or never limiting division.

**FIG. 3.**
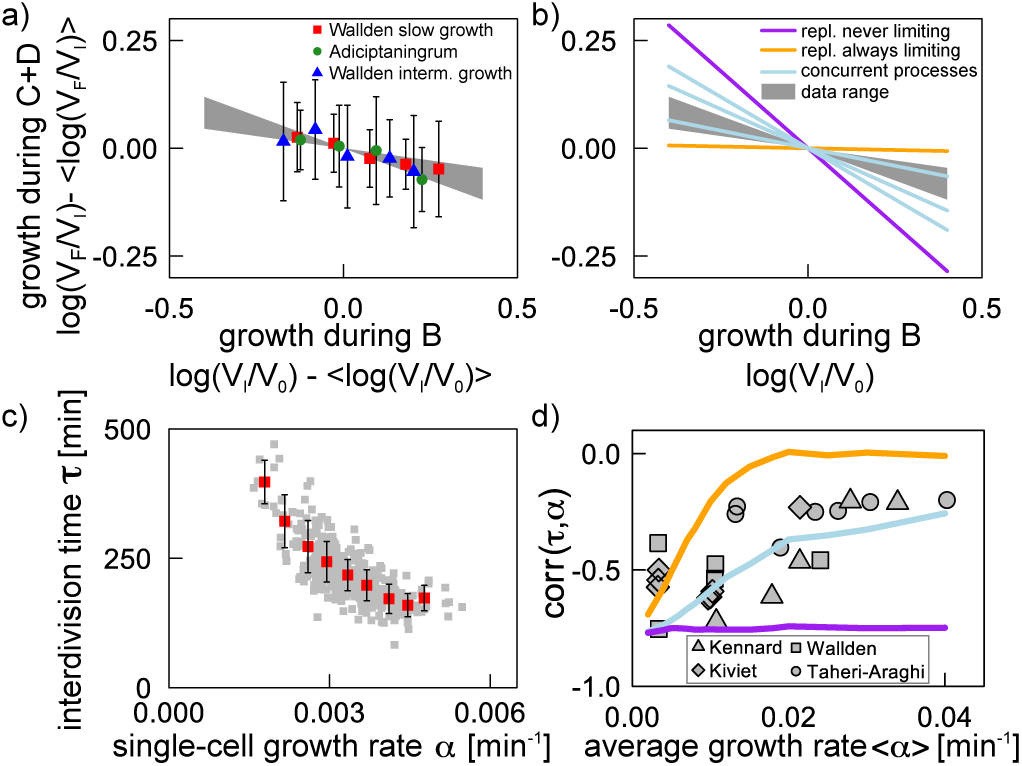
Predictions of the concurrent-processes model beyond the *C*+*D* period are verified in data. a) In data, growth quantified by logarithmic final to initial size ratio) in the *C* + *D* period has a weak negative correlation with growth in the *B* period (see ref. [13]). b) This correlation falls in the range where the replication and inter-division processes compete to set cell division (cyan lines, obtained with same parameters as Fig. 2bd, see methods). c) and d) The negative correlation of inter-division time with individual-cell growth rate (exemplified for one data set in panel c) has a decreasing trend with increasing growth rate, captured by the model (cyan line *p_H_* = 0.25, and other parameters fixed as in the other figures, see methods). Cell-cycle subperiods data from refs [6, 11]. Inter-Division cycle data (in panel d) from refs. [6, 10, 14, 15]

In *E*. *coli*, the average cell size varies across conditions, but the average size per replication origin at initiation remains nearly constant [16]. This old observation was recently proven to be very robust to perturbations [17, 18]. Classically, this observation was interpreted as evidence for a replication-limited cell-division circuit [4, 16]. In a scenario of concurrent processes, size per origin can be constant if the characteristic division size is informed of the characteristic initiation size[19]. Causation could act in either (or both) directions, i.e., the characteristic cell size could be caused by the encoded initiation size per origin, or vice versa (or be the result of a mutual coupling) [20, 21]. In this regard, it is interesting to note that deletion of the nucleoid occlusion SlmA, which prevents division in presence of unsegregated chromosomes, leaves mean cell size unaffected [5, 22], in line with the idea that the replication-associated process does not always drive cell division. The exact causal (and temporal) chain of events might be detectable looking at single cells under perturbations and during nutrient shift experiments. Finally, if we abandon the long-standing assumption that replication initiation always drives cell division in *E*. *coli* favoring a scenario of concurrent processes, we are forced to rethink the regulatory process linking replication to cell division [20], and alternative hypotheses should be revisited in data [23, 24].

More in general, following the pioneering views of Boye and Norstrom [20], our findings support a change of perspective on the cell cycle. We can draw a comparison with the way cell cycle is described in animal cells. A prevalent narrative of the cell cycle, particularly for (higher) eukaryotes, is a succession of well-determined transitions under a common master clock. However, this paradigm struggles to find the “final trigger” of cell-cycle intervals [25]. Instead, in line with our concurrent-processes view, emerging evidence supports the idea that a key aspect could be the integration of *multiple* decisions [25, 26]. For example cell division in higher eukaryotes requires coordination of the actin cortex and the spindle [26], with a necessary crosstalk between these two systems that is analogous to the one addressed here. The concept of concurrent processes on competing time scales can be important to characterize other cell-cycle stages than division, as well as the cell cycles of other organisms. We expect that this different perspective will affect future analyses of the coordination of cell division and other cell-cycle stages with metabolic and physiological cues.

## Author Contributions

MCL MO GM and JG designed research and contributed with key ideas and analyses at different stages. GM relentlessly performed data analysis, model simulations, and analytical calculations; JG provided a forceful drive for the analytical calculations. MCL conceived the project and wrote the paper, with constant help and feedback from the other authors.

## Acknowledgments

We are very grateful to Nancy Kleckner, Clotilde Cadart, Sven van Teeffelen, Ariel Amir, Bianca Sclavi, Sander Tans, Ilaria Iuliani for their feedback. This work was supported by the International Human Frontier Science Program Organization, grant HFSP RGY0070/2014. MO was supported by the “Departments of Excellence 2018 - 2022” Grant awarded by the Italian Ministry of Education, University and Research (MIUR) (L. 232/2016). JG was supported by an Omidyar Postdoctoral Fellowship at the Santa Fe Institute. GM was supported by grant nr. 31003A_169978 from the Swiss National Science Foundation to Martin Ackermann.

## METHODS

### Data analysis

Figs. 2 and 3ac use data on SeqA foci formation from refs [6, 11]. The data contain information on cell division, size vs time, and replication initiation time of single tracked cells. Fig. 3d also uses data sets from refs. [6, 10, 14, 15] The data contain information on cell division, size vs time of single tracked cells for a total of about 10^6^ cells. The dataset in ref. [6] contains about 400 cells in the slow growth condition and about 90 pairs of mother-daughter cells in the intermediate growth condition. The dataset in ref. [11] contains 80 cells. For the slow growth data in ref. [6] and for ref. [11], cells typically have a single DNA replication round during a cell division cycle, and mother-daughter tracking is not necessary. For the intermediate growth conditions of ref. [6] cells typically have two overlapping rounds of DNA replication and mother-daughter tracking is needed to define the *C* + *D* period, since it spans two cell cycles. The plots in Fig. 2 and 3 show binned averages (where the *x*–axis variable was subdivided into bins of equal size); the averages of the *y*–axis variables were considered only for bins containing more than 5 data points.

### Mathematical model and analytical predictions

We describe here the mathematical framework for the concurrent-processes model and the main analytical results. To quantify the coupling between size and cell-cyle progression (and growth) for each interval *X*, the model uses the parameters λ̃_*x*_. Such parameters range from 0 (size threshold) to 1 (timer, no size control). In data, λ̃_*x*_ quantifies the slope of the plot of final vs initial logarithmic size during the cell-cycle interval (see Fig 2ac, where λ̃_*C*+*D*_ is one minus the slope of the so-called “size-growth” plot [27] of relative growth during a cell-cycle interval *vs* the cell size at the entry of the interval). In the model, initiation of replication is set by a size-coupled process acting between consecutive initiations, with a coupling parameter λ̃_*I*_. When λ̃_*I*_ = 1/2, initiation follows an adder [7], while for λ̃_*I*_ = 0, initiation is triggered by a critical size [6]. The case of overlapping replication rounds of DNA replication [28] is described by this model by the assumption that the initiation circuit encodes a size in units of replication origins [7]. Equally, division size is set by a process of strength λ̃_*H*_, which is an adder when λ̃_*H*_ = 1/2. Stochasticity parameters describe size-independent cell-to-cell variability in duration of the inter-division and inter-initiation processes. The “*C* + *D*′” period duration is assumed to be a size-independent Gaussian random variable with assigned mean and variance (a “timer”). Finally, cells are assumed to grow exponentially, and the growth rate is a random variable with prescribed distribution (e.g. Gaussian).

Importantly, the qualitative predictions of the model are very robust to variations in the actual mechanisms of cell-cycle progression, e.g., whether initiation is controlled by a sizer per origin or an adder per origin, or something else, specified by the parameters λ̃_*I*_ and λ̃_*H*_ in the model. This point can be shown by direct analytical estimates. Specifically, the defining equations of the model are

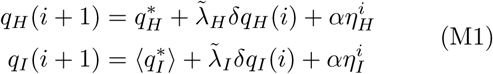

where *i* is a cell cycle index, *q_x_* = *log*(*V_x_*) are logarithmic sizes and *δ_qx_* = *q_x_* – 〈*q_x_*〉 are fluctuations. Finally, *ηx* are noises describing cell-to-cell variability. Eq. (M1) states that size at division and size initiation are corrected with a strength that is specified by the parameters λ̃_*I*_ and λ̃_*H*_ [29].

The logarithm of the final size encoded by the replication-related process in a given cell cycle *i*, *q_R_*, can be expressed as

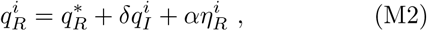

where

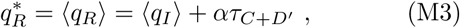

and *τ*_*C*+*D*′_ is the time for the replication-related process to be ready for cell division.

The concurrent processes conditions states that the final size is the one dictated by the slowest process, hence

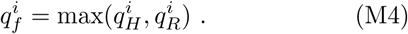

The parameter λ̃_*C*+*D*_ is not an active control in this model, but a result of the interplay between the concurrent processes. It is measured by the slope of Fig. 2ac, or equivalently by the correlation function

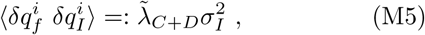

In order to compute this average, we rewrite Eq. (M4) as

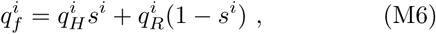

where *s^i^* is a random variable with values in {0,1}. The probability that *s^i^* is 1 depends on *q*_0_ and *q_I_*. We consider the approximation where *s^i^* = 1 with probability *p_H_* independently of *q*_0_ and *q_I_* (but dependently on their averages 〈*q_0_*〉 and 〈*q_I_*〉 and noises, see below). We verified that this approximation works very well with simulations in the noise range of empirical data. *p_H_* is the probability that the inter-division process is rate-limiting, a relevant outcome of the model (depending on the parameters), and is estimated below.

Combining Eqs. (M5), (M6) and (M1), we obtain with some algebra the equation

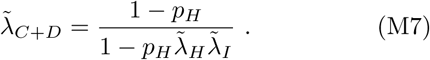

The ratio in the r.h.s of Eq. (M7) is strictly 1 when *p_H_* = 0 (i.e. replication is always rate-limiting), and strictly 0 when *p_H_* = 1 (replication is never limiting). Instead, intermediate values of this ratio can only be attained for intermediate values of *p_H_*, for many combinations of the other parameters. Note that Eq. (M7), given λ̃_*H*_ and λ̃_*I*_, allows to estimate directly *p_H_*. In data, since we can assume that λ̃_*H*_ ≈ 1/2 we estimate that *p_H_* ≈ 0.24–0.69, for a sizer at initiation (λ̃_I_ ≈ 0) and *p_H_* ≈ 0.3 – 0.75 for an adder between initiations (λ̃_*I*_ ≈ 1/2).

Let us now consider the pattern in Fig. 2b. When replication sets division (with probability 1 – *p_H_*), the mean initiation volume 〈*V_I_*〉 dictates the duration of the *C* + *D* period (which corresponds to the *C* + *D*′ period) independently on the growth rate; when instead replication is not the bottleneck (with probability *p_H_*), the final size will be initiation-independent and encoded by the inter-division process 〈*V_H_*〉 and for a fixed growth rate α the duration of the *C* + *D* period will be proportional to (1/*α*)(〈*q_H_*〉 – 〈*q_I_*〉 =: *x*/*α*, where *χ* is roughly the log-ratio of the sizes encoded by the two processes (assuming that log(〈*V_X_*〉) ≈ 〈*q_x_*〉, which is valid for small noise [7, 29]). We can then estimate the effective duration of the *C* + *D* periods as

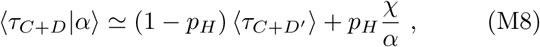

Eq. (M8) is approximate since at fixed single-cell growth rate the probability that the inter-division sets division varies compared to *p_H_* (which is its average value).

Eq. (M8) predicts that the duration of the period is a sum between a constant and an inverse-relationship with the single-cell growth rate. The assumption that replication limits division can only predict the constant part, at odds with data. Conversely, Fig. 2 shows how data are in line with the predictions of the concurrent processes model.

Eq. (M8) contains 〈*q_H_*〉 and 〈*τ*_*C*+*D*′_〉, neither of which is directly accessible from experiments. We can turn it in an expression containing the final (log) size 〈*q_f_*〉, which is measured directly, using the fact that, under the same assumptions,

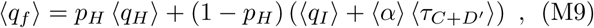

where we have used independence of *C* + *D*′ duration. Substituting this expression in Eq. (M8), we obtain that

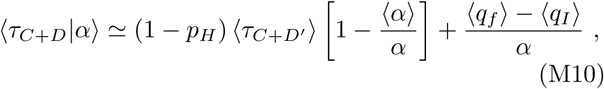

which can be solved for the only unknown parameter, the product (1 – *p_H_*) 〈*τ*_*c*+*D*′_〉 to estimate its value from data, leaving no adjustable parameters for the predictions shown in Fig. 2d.

### Model parameters

We discuss here the values of the model parameters used in the plots, and how they were fixed. The cyan lines in Fig. 2c represent theoretical predictions, and have a negative slope corresponding to Eq. (M7) with *p_H_* = 0.48, *p_H_* = 0.3 and *p_H_* = 0.75 for an adder between initiations (see below), respectively for the Wallden slow growth, Adiciptaningrum, and Wallden intermediate growth data sets [6, 11]. These values were fixed from Fig. 2a and Eq. (M7). The same values of *p_H_* are used in all the other plots. In Fig. 2d, the orange lines are the best fit for the case where replication is limiting (corresponding to the model of ref. [7]). The average duration of the *C* + *D* period is set from the empirical averages to to 160, 102 and 75 minutes, respectively for the Wallden et al. slow growth, Adiciptaningrum et al., and Wallden et al. intermediate growth datasets. Cyan curves follow Eq. (M8) with *p_H_* fixed as above and 〈*τ*_*C*+*D*′_〉 was fixed from Eq. (M10) to the following values: 49 min (Wallden *et al*. slow growth), 52 min (Adiciptaningrum *et al*.), 41 min (Wallden *et al*. intermediate growth), respectively.

Fig. 3bd show the result of numerical simulations, where the parameters were constrained from data. The plots in Fig. 2 fix the values of *p_H_* and 〈*τ*_*C*+*D*′_〉 for the three data sets in Fig. 3b, while Fig. 3d corresponds to a wide range of growth conditions and required more general choices. Specifically, the numerical simulations use the following input parameters, directly fixed from data: (i, ii) The average growth rate 〈*α*〉 and its variance
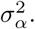
In Fig. 3d, the average growth rates range from 0.002 to 0.04 min^−1^ to match the experimental values. The variance of the growth rate was chosen by keeping a constant CV of 0.15. The growth rate was a normal or log-normal random variable, (the two choices do not affect the results, see below), extracted independently for every cell cycle. (iii, iv) The average added volume per origin between consecutive initiations 〈*v_I_*〉 and its variance
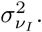
The average added volume per origins was fixed to 0.45 *μ*m^3^ as measured in ref. [6], and the variance was fixed to a constant CV of 0.15. The added volume between consecutive initiations was assumed to be log-normally distributed (and extracted independently for every cell cycle). (v, vi) The average time needed for replication and segregation (the *C* + *D*′ period) 〈*τ*_*C*+*D′*_〉 and its variance
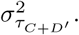
In Fig. 3d, The average duration of the *C* + *D*′ period was set to 45 min independently from the average growth rate, and its CV was maintained constant and equal to 0.2 for the simulations corresponding to replication always limiting (orange lines) and concurrent processes (cyan lines). The average duration of the *C* + *D*′ period was set to 0 for the simulations where replication is never limiting (purple lines). *τ*_*C*+*D*′_ was assumed to be a normal random variable (extracted independently for every cell cycle). Note that the average added volume per origin between consecutive initiations, the average growth rate and the average *C* + *D*′ duration set the characteristic division size of the inter-initiation process,
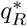
≈〈log (*v_I_*)〉 + 〈*τ*_*C*+*D*′_〉 〈*α*〉. (vii, viii) The average inter-division added volume 〈Δ_*H*_〉 and its variance
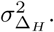
Note that 〈Δ_*H*_〉 sets the characteristic size of the inter-division process
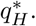
In Fig. 3b, 〈Δ_*H*_〉 was fixed to achieve the same values of *p_H_* as in Fig. 2bd. In Fig. 3d, the inter-division added volume was fixed so that for the case of concurrent cycles the replication-related process and the inter-division process would compete to set division in the same way across all conditions. Specifically, we chose 〈Δ_*H*_〉 = 〈*v_i_*〉 exp (45min o 〈*α*〉) for the case of concurrent processes (cyan line in Fig. 3d). The case where replication is never limiting (purple line) was simulated by assuming the same 〈Δ_*H*_〉 but setting *τ*_*C*+*D*′_ to zero. Finally, we set 〈Δ_*H*_〉 = 0 to simulate the case of replication always limiting (orange line). With these choices, *p_H_* ≈ 0 for the replication always limiting case (orange line in Fig. 3d), *p_H_* ≈ 0.6 for concurrent cycles (cyan line) and *p_H_* ≈ 1 for the replication never limiting models (purple line). The variance
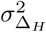
was set to a constant CV of 0.15 over the different growth conditions. The inter-division volume was assumed to be a log-normally distributed random variable as above.

We chose to present data with λ̃_*I*_ = λ̃_*H*_ = 1/2 (adder) simulations, but we explored other values of these parameters, and in particular λ̃_*I*_ = 0, λ̃_*H*_ = 1/2, initiation triggered at fixed size per origin, finding very robust results (data not shown). Additionally, while Fig. 3 follows a specific set of parameters, we explored systematically in simulations different characteristic sizes and noise levels for the two processes. For instance, as mentioned above, the simulations for Gaussian distributed growth rate, inter-division volume, and added volume per origins between consecutive initiations show no substantial differences in the trends shown in Fig. 3 (data not shown).

